# Diversity and distribution of sulfur metabolism in the human gut microbiome and its association with colorectal cancer

**DOI:** 10.1101/2021.07.01.450790

**Authors:** Patricia G. Wolf, Elise S. Cowley, Adam Breister, Sarah Matatov, Luke Lucio, Paige Polak, Jason M. Ridlon, H. Rex Gaskins, Karthik Anantharaman

**Affiliations:** Institute for Health Research and Policy, University of Illinois at Chicago, Chicago, IL, USA; University of Illinois Cancer Center, University of Illinois at Chicago, Chicago, IL, USA; Department of Animal Sciences, University of Illinois Urbana-Champaign, Urbana, IL, USA; Division of Nutritional Sciences, University of Illinois Urbana-Champaign, Urbana, IL, USA; Department of Bacteriology, University of Wisconsin-Madison, Madison, WI, USA; Microbiology Doctoral Training Program, University of Wisconsin-Madison, Madison, WI, USA; Carl R. Woese Institute for Genomic Biology, University of Illinois Urbana-Champaign, Urbana, IL, USA; Cancer Center at Illinois, University of Illinois Urbana-Champaign, Urbana, IL, USA; Department of Biomedical and Translational Sciences, University of Illinois Urbana-Champaign, Urbana, IL, USA; Department of Pathobiology, University of Illinois Urbana-Champaign, Urbana, IL, USA

## Abstract

Microbial sulfidogenesis produces genotoxic hydrogen sulfide (H_2_S) in the human gut using inorganic (sulfate) and organic (taurine/cysteine/methionine) substrates, however the majority of studies have focused on sulfate reduction using dissimilatory sulfite reductases (Dsr). Recent evidence implicates microbial sulfidogenesis as a potential trigger of colorectal cancer (CRC), highlighting the need for comprehensive knowledge of sulfur metabolism within the human gut. Here we show that microbial sulfur metabolism is more abundant and diverse than previously studied and is statistically associated with CRC. Using ~17,000 bacterial genomes from publicly available stool metagenomes, we studied the diversity of sulfur metabolic genes in 667 participants across different health statuses: healthy, adenoma, and carcinoma. Sulfidogenic genes were harbored by 142 bacterial genera and both organic and inorganic sulfidogenic genes were associated with carcinoma. Significantly, anaerobic sulfite reductases were twice as abundant as *dsr*. We identified twelve potential pathways for reductive taurine metabolism including novel pathways, and prevalence of organic sulfur metabolic genes indicate these substrates may be the most abundant source of microbially derived H_2_S. Our findings significantly expand knowledge of microbial sulfur metabolism in the human gut, and highlight key gaps that limit understanding of its potential contributions to the pathogenesis of CRC.

## INTRODUCTION

The human gut is a dynamic nutrient rich environment that harbors a diverse metabolically active microbial community. It has been established that human health and disease are inextricably linked to microbial composition, however much remains unknown regarding the functional capacity of human gut microbes^1–3^. This has manifested in bacteria or their niches being loosely characterized as “beneficial”, “commensal”, or “deleterious”, which is problematic as microbial functionality is often species-specific and microbes are capable of metabolic shifts based on available substrates^4,5^. Genomic approaches have enabled rapid discovery of novel bacteria whose functional characteristics have yet to be characterized^6^. These discoveries have filled gaps in knowledge regarding the metabolic capacity of human gut microbes and allow design of hypothesis driven interventions that create beneficial shifts in microbial communities. This approach may have particularly important implications in human diseases for which associations between microbial composition, dietary intake, and disease risks have been established.

For example, there is strong evidence linking a diet high in red and processed meat with colorectal cancer (CRC)^7^. In addition, bacteria capable of producing hydrogen sulfide (H_2_S) are associated with a western diet^8,9^, colonic inflammation^10^, and CRC^11–19^. At μM concentrations^20^, endogenously produced H_2_S can act as a vasorelaxant^21^, reduce endoplasmic reticulum stress^22^, and prevent apoptosis^23^. At millimolar concentrations, as commonly found in the colon, H_2_S inhibits cytochrome oxidase causing reductive stress and is genotoxic^24–27^. Previous work investigating microbial sulfidogenesis in the human gut have mostly focused on sulfate reducing bacteria (SRB) that perform inorganic sulfur metabolism^28^. However, recent evidence indicates that organic sulfur metabolism by gut bacteria may be a key mechanism linking diet and CRC^29^. Indeed, CRC-associated bacteria have been shown to produce H_2_S via metabolism of sulfur amino acids^6^, and the taurine metabolizing *Bilophila wadsworthia* was found previously to be a significant indicator of CRC^11^. Consumption of a diet high in red and processed meat increases colonic concentrations of organic sulfur, which may increase colonic concentrations of microbially derived H_2_S to genotoxic levels^11,30,31^. This suggests that sulfur metabolism in the human gut microbiome may be more widespread than originally believed, and exposes current gaps in our knowledge of the metabolic functions of CRC-associated bacteria.

Thus, the objective of this study was to use genomic and metagenomic tools to gain a greater understanding of the sulfidogenic capacity of the human gut microbiome. To do so, we investigated the prevalence of sulfidogenic genes in gastrointestinal bacterial genomes, established a network of sulfur metabolic transformations, and identified novel sulfidogenic bacteria. Using newly developed gene databases, 16,936 publicly available bacterial metagenome assembled genomes (MAGs) from human gut microbiomes were mined to compare the relative contribution of inorganic and organic sulfidogenic genes to microbial sulfur metabolism. Gene presence was then compared among disease states in five CRC microbiome studies to evaluate potential contributions of microbial H_2_S production to CRC risk. This study provides the most comprehensive analysis of microbial sulfur metabolism in the human gut to date, and thereby provides a platform for hypothesis-driven experiments characterizing the role of sulfur metabolites in CRC and other inflammatory-associated gut disorders.

## RESULTS

### Common pathways of microbial sulfur metabolism are prevalent in human gut microbiomes

To understand the diversity, distribution, and ecology of microbial sulfur metabolism in the human gut and its implications in disease, we investigated the complex pathways for sulfur transformations (Fig. 1). Stool shotgun metagenomic sequence data were used from 5 publicly available studies that investigated the gut microbiome in healthy subjects and patients who had adenoma or carcinoma of the colon. Collectively, 265 healthy participants, 112 participants with adenoma, and 290 participants with carcinoma were examined. Participant location, associated metadata, and study references are listed in (Table 1)^17,32–35^. A total of 16,936 bacterial metagenome assembled genomes (MAGs) were recovered from a previous study that used standardized bioinformatic pipelines for metagenome assembly^36^. A concatenated ribosomal protein tree was created to examine the full diversity of the samples used in this study with the majority of genomes being classified in the phyla Firmicutes, Proteobacteria, and Actinobacteria (Fig. S1). Open reading frames (ORFs) were then predicted followed by homology-based identification of 72 genes associated with microbial sulfur metabolism (Table S1) using available and custom Hidden Markov Models (HMMs). These analyses enabled us to determine the breadth of microbial sulfur metabolism in the human gut and its associations with CRC (Table S2, Fig. S1).

**Figure 1.**
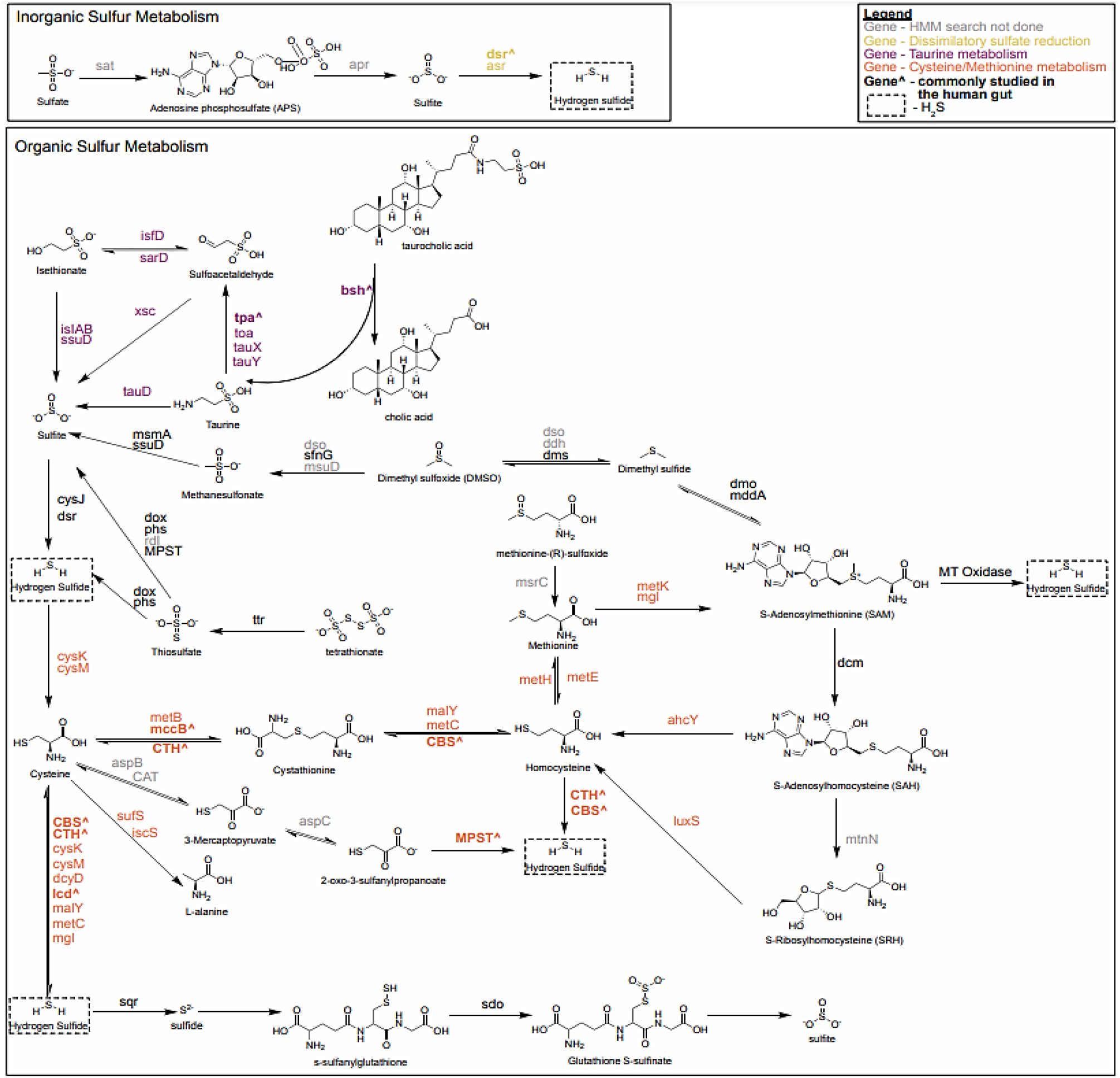
Potential microbial sulfur transformations in the human gut microbiome. Microbial sulfur metabolism results in the production of genotoxic H_2_S (dashed box) via metabolism of inorganic sulfate (yellow) or organic sulfur amino acids like cysteine and methionine (maroon), or taurine (orange). Previous studies of microbial sulfidogenesis in the human gut have focused mainly on genes harbored by Bilophila, Fusobacterium, and the sulfate reducing bacteria (bolded with a “^”). All genes listed were analyzed in this study except those listed in gray. Reactions are not balanced and only the main sulfur component reactants and products are shown. Some intermediate steps are not shown.

**Table 1.** Overview of original datasets used for this study.

| Study | Country of Participant Recruitment | Disease State | Number of Participants | Number of MAGs |
| --- | --- | --- | --- | --- |
| Feng<sup>22</sup> | Austria | Control/Healthy | 61 | 1690 |
|  |  | Adenoma | 47 | 1421 |
|  |  | Carcinoma | 46 | 1418 |
| Hannigan<sup>23</sup> | United States, Canada | Control/Healthy | 26 | 101 |
|  |  | Adenoma | 23 | 57 |
|  |  | Carcinoma | 26 | 78 |
| Vogtmann<sup>24</sup> | United States | Control/Healthy | 58 | 1863 |
|  |  | Adenoma | 0 | 0 |
|  |  | Carcinoma | 52 | 1622 |
| Yu<sup>25</sup> | China | Control/Healthy | 54 | 1397 |
|  |  | Adenoma | 0 | 0 |
|  |  | Carcinoma | 75 | 1910 |
| Zeller<sup>17</sup> | France | Control/Healthy | 66 | 1839 |
|  |  | Adenoma | 42 | 956 |
|  |  | Carcinoma | 91 | 2584 |
| Totals |  | Control/Healthy | 265 | 6890 |
|  |  | Adenoma | 112 | 2434 |
|  |  | Carcinoma | 290 | 7612 |

To understand the prevalence of microbial sulfur metabolism in a representative human gut microbiome, 514 gastrointestinal microbial genomes were obtained from the Human Microbiome Project (HMP) and surveyed for common metabolic pathways associated with H_2_S production from cysteine, taurine, and sulfate/sulfite. Functional genes encoding proteins for cysteine metabolism made up the majority of sequences and included cystathionine-β-lyase (*malY, metC*) (11.8%), cystathionine-β-synthase (*CBS*) (27.5%), cysteine desulfhydrase (*lcd*) (2.6%), D-cysteine desulfhydrase (*dcyD*) (17.9%), and methionine-γ-lyase (*mgl*) (14.7%). In contrast, dissimilatory sulfite reductases (*dsrAB*) catalyzing the final step of sulfate and taurine respiration to form H_2_S, made up only 18.2% of identified genes (Fig. S2). In total, 313 sulfidogenic genes were identified from 183 bacterial genomes (35.6% of total genomes), and spanned across six phyla including Proteobacteria (36.1%), Firmicutes (26.2%), Bacteroidetes (20.2%), Fusobacteria (15.8%), Actinobacteria (1.1%), and Synergistetes (0.5%). This key finding demonstrates that pathways for sulfur metabolism were prevalent in human gut microbiome genomes and that cysteine may be an underestimated substrate for microbial sulfur metabolism in the gut.

### Genes for anaerobic sulfite reductases (*asrABC*) are more prevalent than dissimilatory sulfite reductases (*dsrAB*) in the human gut

Of the limited literature regarding human colonic sulfidogenic bacteria, the most well studied are the SRB who are capable of reducing inorganic sulfate supplied exogenously by diet or endogenously by degradation of sulfated bile acids and mucins (estimated 1.5-16 and 0.96-2.6 mmol/day respectively)^28,37–40^. Two enzymes are able to complete the final step of the reaction which catalyzes the six-electron reduction of sulfite to H_2_S — dissimilatory sulfite reductase (Dsr) and anaerobic sulfite reductase (Asr) (Fig. 1). It has been proposed that sulfite reduction takes place as a series of two electron transfers to DsrAB from the DsrMKJOP complex via DsrC^41^. Genes for the Dsr pathway, *dsrAB*, are highly conserved among SRB and diversely distributed among phyla in environmental samples^42^. However, culture and PCR based studies of SRB diversity in human stool and colonic mucosa indicate that *dsrAB* is harbored by only five resident genera namely *Bilophila* spp., *Desulfovibrio* spp., *Desulfobulbus* spp., *Desulfobacter* spp., and *Desulfotomaculum* spp^43–46^.

Within our database of 16,936 bacterial MAGs from human gut samples, *dsrAB* genes were present in 121 MAGs (<1% of total MAGs), and in 17.4% of total subjects (Table S3.). Taxonomic classification demonstrated genera commonly associated with Dsr activity in the gut microbiome were represented including *Desulfovibrio* spp. and *Bilophila* spp. In addition, our results revealed six genera not commonly ascribed as human gut SRBs namely *Colinsella* spp., *Eggerthella* spp., *Enterococcus* spp., *Flavinofracter* spp., *Gordonibacter* spp., and *Roseburia* spp (Tables S2, S4). Since previous phylogenetic analyses in environmental samples indicated that *dsrAB* acquisition was often the result of multiple lateral gene transfer events^42^, a concatenated gene tree was generated with a reference database of *dsrAB* sequences to observe the consensus phylogeny of *dsrAB* sequences in human gut bacteria. *DsrAB* sequences from sample MAGs separated into three distinct clusters which corresponded with the phylogeny of the representative genera: Actinobacteria, Firmicutes, and Proteobacteria (Fig. 2A). This suggests that lateral gene transfer of *dsrAB* genes may be less common in SRBs of the human gut than those observed in environmental studies^42^.

**Figure 2.**
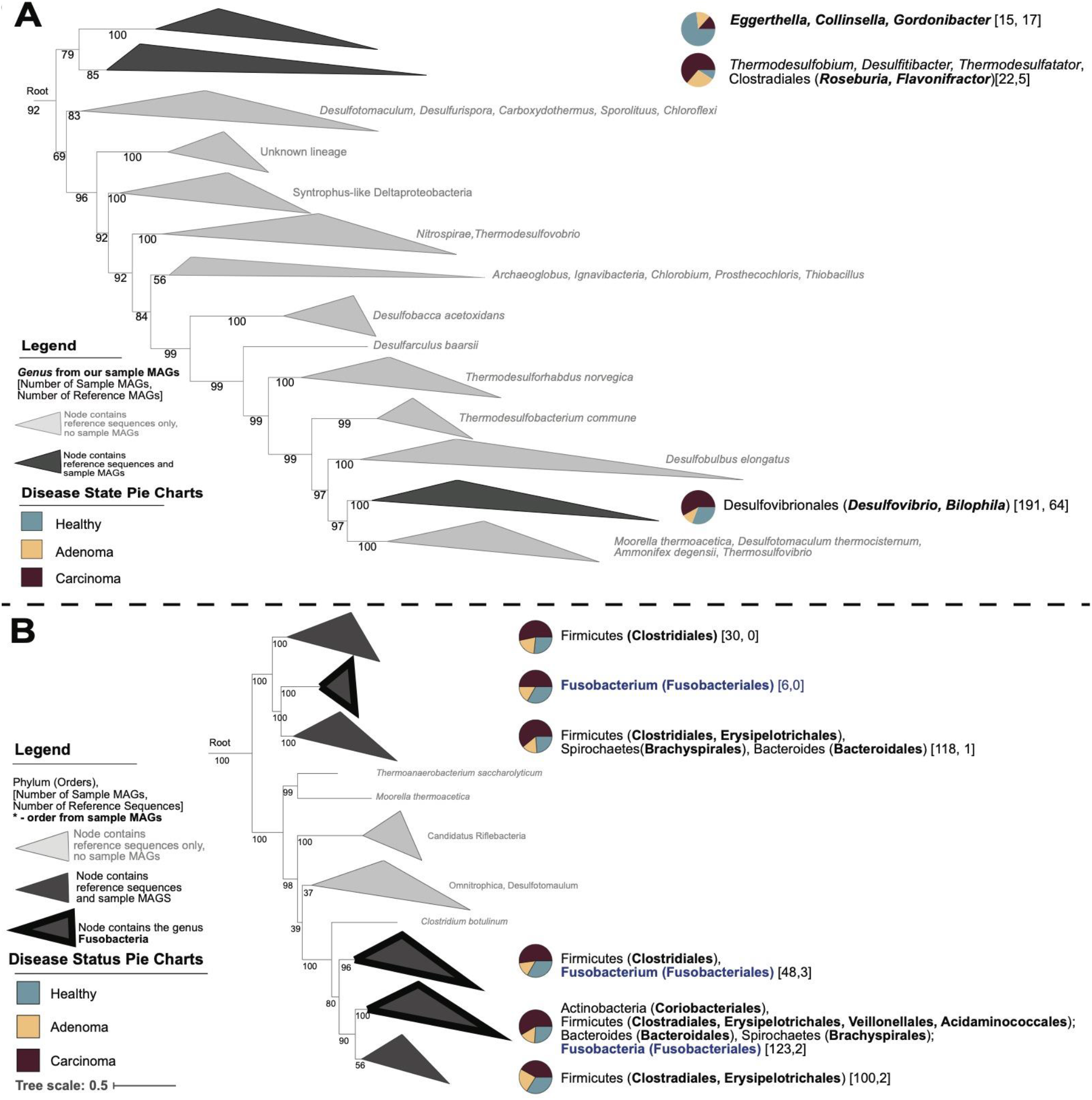
Concatenated protein trees for dissimilatory sulfate reduction pathways. A. Concatenated protein tree showing the diversity of bacteria that possess genes for the final enzyme of the dissimilatory sulfate reduction pathway — dsrAB. B. Concatenated protein tree showing the diversity of bacteria that possess genes for anaerobic sulfite reductase (asrABC), an enzyme also capable of dissimilatory sulfate reduction. Gray clades only contain reference sequences, darker gray clades contain reference sequences and sequences from this study. Bracketed numbers indicate the sequence origin within each clade: [number of sequences from our study, number of sequences from references]. Bacterial genera (dsr) or orders (asr) originating from study samples are bolded. Pie charts indicate the disease state associated with sequences within each clade with blue indicating healthy, yellow adenoma, and maroon carcinoma. Clades outlined in black contain Fusobacterium sequences.

While SRBs may be the most well studied sulfur metabolizing bacteria of the human gut microbiota, studies have focused mainly on bacteria harboring Dsr enzymes. Similar to Dsr enzymes, Asr performs a 6-electron reduction of sulfite to H_2_S (Fig. 1). A recent analysis of gut microbiome including *Fusobacterium nucleatum* and *Clostridium intestinale*^47^. To determine if gut bacteria harbor *asrABC*, we searched the MAG database, revealing that *asrABC* genes were more prevalent in all MAGs and participants than *dsrAB* genes. The *asrABC* genes were present in approximately 2% of total MAGs (388, 390, and 375 MAGs, respectively), and approximately 35% of total subjects, (Table S3). Intriguingly, less than a quarter of the subjects that harbored *asrABC* genes also possessed genes for *dsrAB* (53 of 239 subjects). Taxonomic assignments showed thirty-one genera possessed *asrABC*, spread among five phyla including Actinobacteria, Firmicutes, Fusobacteria, Spirochaetes, and a phylum not previously shown to possess *asrABC* — Bacteroidetes^47^. A consensus tree of reference and sample MAG concatenated *asrABC* sequences revealed clustering of *asrABC* genes across six nodes that were distinct from sample MAG phylogeny. Notably, *asrABC* genes possessed by *Fusobacterium* spp. did not cluster together, but were observed in three separate nodes suggesting this pathway was acquired via multiple lateral gene transfers (Fig. 2B). Together, these data indicate that Asr enzymes may be more important contributors to sulfate and sulfite reduction than Dsr in the human gut, and that bacteria acquired these enzymes via a divergent phylogenetic history.

### Diverse bacteria harbor pathways for taurine metabolism

Sulfonates are organic sulfur compounds with a SO_3_^−^ moiety, that are abundant in marine sediments and detergents, and play an important role in the environmental sulfur cycle^48^. While less is known about the role of sulfonates in the human intestine, there is evidence that metabolism of sulfonates like isethionate and taurine are performed by resident gut microbes^49, 50^. microbial taurine metabolism has gained considerable interest after it was implicated as a potential dietary mechanism of colitis and CRC disparities^10,11^. Representing 18.3% of total free amino acids in the colonic mucosa (13.6 ± 0.5 mmol/kg), taurine is the second most abundant free amino acid in this tissue^51^. Taurine is provided as a substrate to the human gut microbiome either directly through diet or through hydrolysis of taurine conjugated bile acids by the enzyme bile salt hydrolase (BSH). Excess consumption of taurine and cysteine increases tauro-conjugation of secreted bile acids, thus potentially providing additional substrates for bacteria with BSH activity^29^. The *bsh* gene was identified to be present in 15% of total MAGs, and in 89% of total subjects (Table S3). This prevalence was unsurprising, as it has been proposed that hydrolysis of conjugated bile acids may be a detoxification strategy to decrease bile acid toxicity^52^, or may serve as a source of nutrients for microbial growth and energy metabolism^28^.

Once liberated via BSH or made available through dietary intake, taurine can be metabolized by gut bacteria via oxidative or reductive pathways. Taurine oxidation using the enzyme taurine dioxygenase (TauD) is often disregarded as a source of H_2_S production in the anaerobic environment of the human colon. However, previous studies have reported an increase of aerotolerant bacteria adherent to the mucosal surface suggesting a luminal gradient of oxygen provided by host tissues^53^. Assessment of *tauD* gene abundance demonstrated that while the enzyme was present in only 1% of total MAGs, these MAGs were present in 23% of subjects (Table S3). As expected, *tauD* genes were present in six facultative genera (*Eschericia* spp., *Enterobacter* spp., *Citrobacter* spp., *Morganella* spp., *Hafnia* spp., and *Raoultella* spp.), however none of these genera possessed genes for anaerobic or dissimilatory sulfite reduction (Table S4). Thus, TauD appears to be primarily used for taurine assimilation and not H_2_S production.

The only known bacterium to possess the reductive pathway of taurine metabolism in the human gut is *B. wadsworthia*. However, given the taurine rich environment of the colon, it is likely that other bacteria capable of performing this metabolism remain to be discovered. Thus, to identify candidates that may have the capacity to produce H_2_S from taurine in an anaerobic environment, HMM searches of the described cohorts were performed targeting pathways as shown in (Fig. 3).

**Figure 3.**
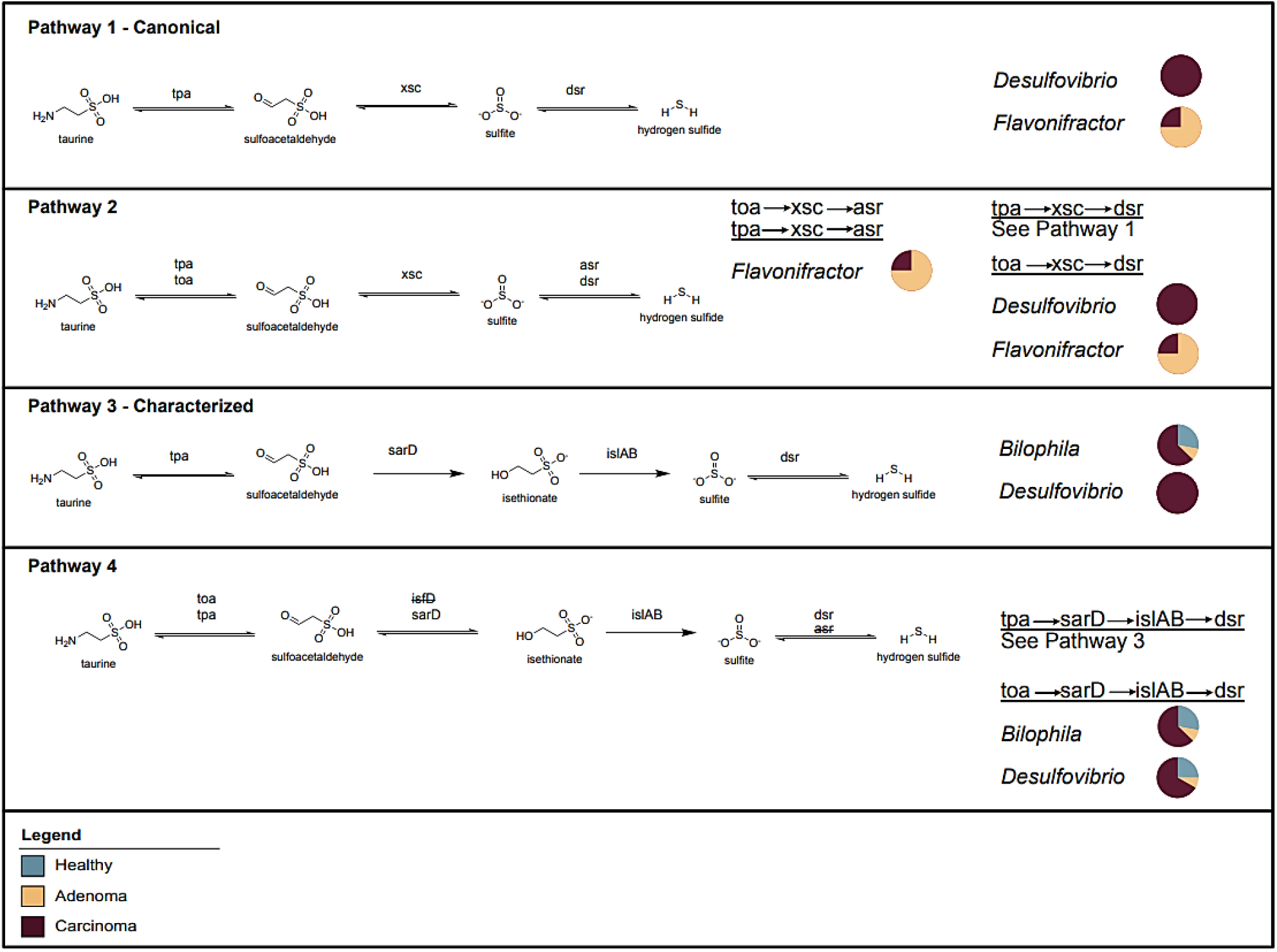
Characterized and proposed pathways of microbial taurine reduction to H_2_S. Pathway 1 - the canonical pathway of taurine reduction in *Bilophila wadsworthia*. Pathway 2 - putative 3-step reactions for taurine reduction analyzed in this study. Pathway 3 - the recently characterized pathway for taurine reduction in *Bilophila wadsworthia*. Pathway 4 - putative 4-step reactions for taurine reduction analyzed in this study. No complete pathways were found involving genes that are struck through. Genera possessing genes for each complete pathway are listed. Only genera listed were found to have complete pathways. Pie *charts* indicate the disease state associated with MAGs of each genus with the specified pathway with blue indicating healthy, yellow indicating adenoma, and maroon indicating carcinoma.

Pathway 1 describes the putative taurine reduction pathway previously thought to be possessed by *B. wadsworthia*^28^. Our search confirmed that this pathway was not possessed by *Bilophila* spp., as recently described^54^. Instead, analyses revealed two genera that harbored genes for this pathway, namely *Desulfovibrio* spp. and *Flavonifractor* spp. The first step of taurine reduction involves the liberation of the nitrogenous group from taurine producing pyruvate or 2-oxoglutarate via the enzymes taurine-pyruvate aminotransferase (Tpa) or taurine-2-oxoglutarate transaminase (Toa), respectively (Pathway 2). For all MAGs that possess the latter 2 genes of this pathway, the genes *tpa* and *toa* co-occur. Notably, *Flavonifractor* spp. also harbors genes for the *asrABC* complex, indicating an alternative final step of the taurine reductive pathway (Pathway 2). Additionally, evaluation of three-step pathway combinations revealed 34 genera that possessed the first and final pathway steps, suggesting the pervasiveness of metabolic cooperation in the gut and also potentially revealing targets for the identification of novel sulfoacetaldehyde acetyltransferases (Xsc) (Fig. 3) (Tables S4, S5).

Pathway 3 represents the recently characterized pathway for taurine reduction in *B. wadsworthia* (Fig. 3)^54^. Gene searches corroborated that *Bilophila* spp. harbored all four genes in this newly defined pathway. However, contrary to what was recently reported, all four genes were also observed in *Desulfovibrio* spp. Since MAGs are unable to allow for granularity at the species level, future work is needed to determine if this pathway is indeed more widespread in resident *Desulfovibrio* species of the human gut. Seven MAGs annotated as *Desulfovibrio* spp. and *Bilophila* spp. also possessed *toa* in the absence of *tpa*, together, this indicates an alternative first step to this pathway (Pathway 4). Notably, unlike the three step Pathway 2, MAGs that possess all genes for the four-step pathway harbor only *dsrAB* and not *asrABC*. In addition, evaluation of four step pathway combinations revealed 7 genera missing either the second or third step of the pathway revealing additional targets for gene discovery (Fig. 3) (Tables S4, S5).

### Cysteine and methionine are understudied and abundant sources of microbially-derived H_2_S in the human gut

Cysteine is a conditionally essential amino acid, which is provided to the intestine directly by diet or the decomposition of methionine (Fig. 1). Methionine restriction alters microbial composition of the gut and down-regulates inflammatory pathways related to oxidative stress^55,56^. In addition, the production of H_2_S via cysteine degradation supports microbial growth and protects from oxidative stress in response to antibiotic treatment^57^. Cysteine metabolizing bacteria are implicated as a source of oral abscess, breath malodor, and delayed wound healing in the oral cavity, and have been repeatedly associated with CRC^12,14,16,17^. Of late, *F. nucleatum* has been of particular interest in CRC^14,16,58^. Studies have demonstrated association between *F. nucleatum* and the tumor surface in a subset of CRC^12,14,59^, *F. nucleatum* DNA in CRC tumors correlate with reduced survival^60,61^, and two subspecies of *F. nucleatum* (*vincentii* and *animalis*) have been proposed as part of a microbial signature for fecal-based CRC classification^17^. However, few studies appreciate that *F. nucleatum* is sulfidogenic, and many other bacteria can also produce H_2_S from cysteine in the human gut.

To gain an appreciation of the abundance of cysteine metabolism within the human microbiome, MAGs were searched for the following genes associated with cysteine metabolism *dcyD, malY, metC, mgl*, as well as cysteine synthase (*cysK, cysM*), cysteine desulfurase (*iscS, sufS*), and cystathionine-γ-synthase (*metB*), and the three human orthologs cystathionine-β-synthase (*CBS*), cystathionine-γ-lyase (*CSE/mccB*), and 3-mercaptopyruvate sulfurtransferase (*3MST*). Genes for upstream pathways of methionine and homocysteine metabolism were also analyzed (Fig. 1, Table S1). Searches revealed that all cysteine metabolizing genes were highly present, with *cysK, lcd, malY*, and *sufS* observed at least once in over 96%of subjects (Table S3). Even those genes that were observed in only 2-5% of total MAGs — namely *dcyD, metC, cysM*, and *mgl* — were still observed to be present in at least 40% of subjects (Table S3). In accordance with this, genes for microbial pathways for methionine metabolism were also highly present in subjects, indicating that methionine may be an important source of microbial derived cysteine in the human gut (Table S3). In addition to being abundantly present, genes for cysteine metabolism were also diversely distributed among 13 phyla and 141 genera. Amongst these, 84 genera have not been previously characterized as sulfidogenic. (Table S4). This may be particularly important in the context of a western diet, as studies using fecal homogenates demonstrate higher production of H_2_S from organic sulfur amino acids compared to inorganic sulfate^62^, and higher protein intake increases ileal output of protein (2.69 vs. 7.45 g/day) and free amino acids (6.90 vs. 20.48 μmol/mL)^63^. Although additional work is needed to comprehensively resolve cysteine metabolism, together these data indicate that the sulfur amino acids cysteine and methionine may be an understudied and abundant source of microbially-derived H_2_S in the human gut.

### Microbial sulfur metabolism is statistically associated with colorectal cancer

Sulfate reducing bacteria and H_2_S have been implicated in CRC pathogenesis^24,25,30,64,65^, however associations of other microbial sulfur metabolism genes with CRC have been less well studied. The proportion of participants in each health status (healthy, adenoma, or carcinoma as evaluated by colonoscopy) category with at least one copy of each gene was determined along with the proportion of the total number of MAGs in each health status with the gene (Fig. 4, Table S3). Genes involved with cysteine and methionine metabolism were generally abundant regardless of health status (Fig. 4A, size of dots) and abundant in many MAGs (Fig. 4A, color of dots). Genes involved in sulfur and taurine metabolism were variable in their distribution among participants in the 3 health states and in MAGs (Fig. 4).

**Figure 4.**
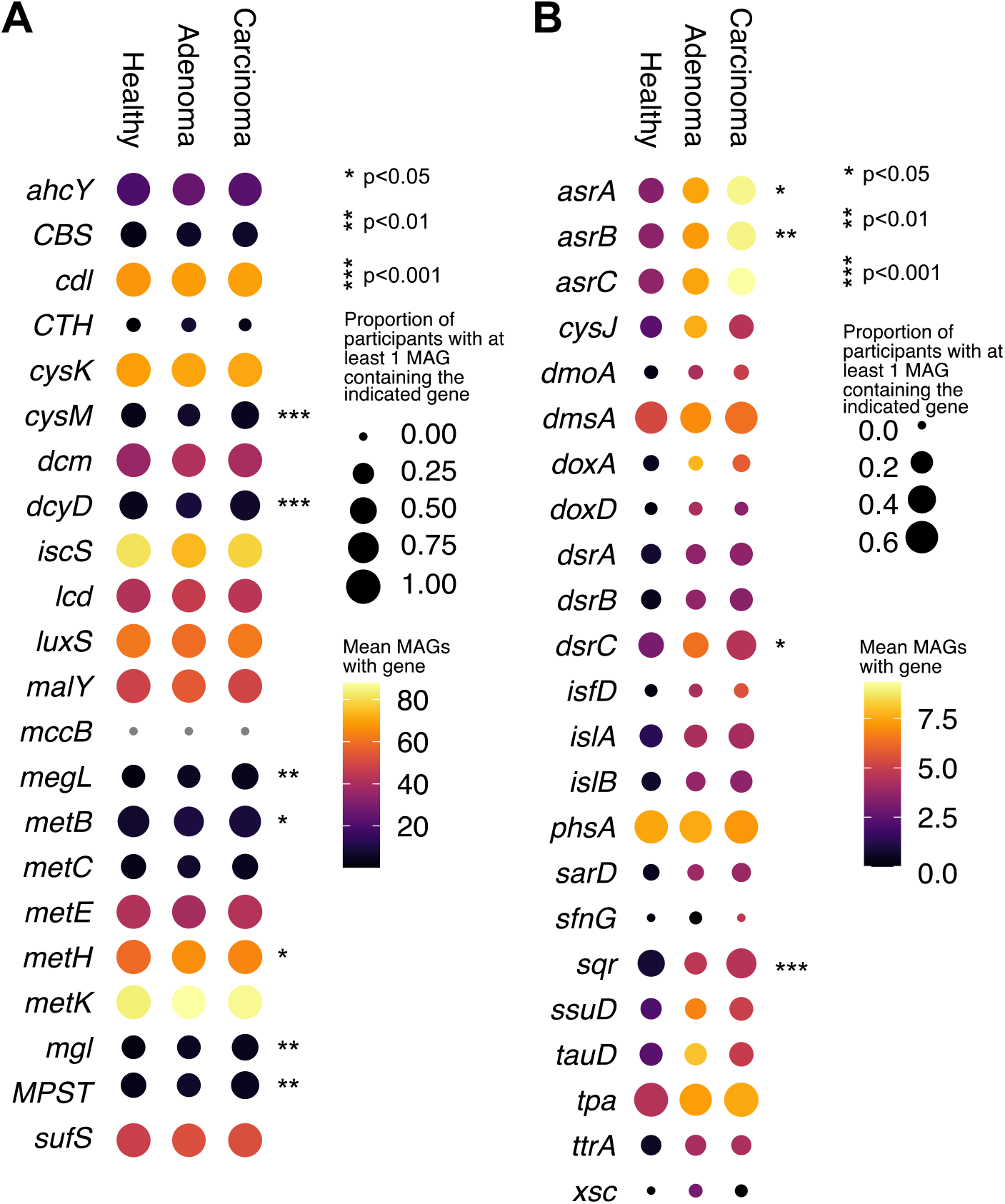
Genes for microbial sulfur metabolism are abundant and significantly associated with colorectal cancer. Dot plots of selected genes related to microbial cysteine and methionine metabolism (A) and taurine and sulfur metabolism (B) across three disease states: healthy, adenoma, and carcinoma. The size of each dot indicates the proportion of participants in each disease state with at least 1 copy of the indicated gene in their bacterial MAGs and the color of each dot indicates the mean number of MAGs with that gene in the subset of participants that have at least 1 copy of the gene. Genes that have a non-random distribution across disease status as analyzed by chi-squared analysis are indicated by asterisks.

To determine if sulfur metabolism was associated with CRC, we conducted statistical tests to study the distribution of sulfur genes in participants across each health status. Genes for tetrathionate metabolism, which was previously implicated as an electron acceptor provided by gut inflammation^66^, were present in less than 1% of MAGs and were not significantly different among the three health statuses. Genes involved in cysteine and methionine metabolism including *cysM, dcyD, mgl, metB, metH*, and *sdo*, were found to have distributions that were significantly different (at least p-value< 0.05) among participants in the 3 health states (Fig. 4A, Table S6, Fig. S5), with *sdo, cysM, mgl, metB*, and *metH* being more likely to be found in carcinoma. While 92 genera with cysteine metabolizing genes have been associated with CRC previously, our results indicate that this association may involve sulfidogenesis. Indeed, MAGs for cysteine metabolizing genes were pervasive in genera most commonly associated with CRC, corroborating recent work that observed that genes for cysteine metabolism were significantly more abundant in subjects with CRC^67^ (Fig. S3). Sulfur and taurine metabolism genes, *asrA, asrB, dsrC*, and *sqr*, had distributions that were significantly different among the 3 health states with all being more likely to be found in carcinoma (Fig. 4B, Table S6, Fig. S4,S5).

Intriguingly, for genes related to dissimilatory sulfate reduction, different organisms exhibited different clustering patterns based on both phylogeny and disease state. For the Dsr pathway, the majority of *Desulfovibrio* spp., *Bilophila* spp., *Flavinofracter* spp., and *Roseburia* spp. were harbored by participants with carcinoma while the majority of *Colinsella* spp., *Eggerthella* spp., and *Gordonibacter* spp. originated from healthy participants (Fig. 2A - pie charts). For the Asr pathway, the majority of sequences were recovered from participants with carcinoma (Fig. 2B - pie charts). These data demonstrate that genes for sulfur metabolism are associated with carcinoma.

To gain a deeper understanding of the ecology of microbial sulfur metabolism during colorectal carcinogenesis, we first determined the associations of sulfidogenic genes between different CRC stages from the three datasets that reported staging data, and then investigated growth rates of selected taxa previously considered to be microbial markers of CRC^11,17^. The likelihood of patients having sulfidogenic genes was not significantly different between stages (Supplemental Table 8). However, participants were more likely to have genes for inorganic sulfur metabolism in earlier stages of CRC, while participants having genes for organic sulfur metabolism was uniformly high (Fig S6). *Mgl* — an organic sulfur metabolizing gene — was increasingly present across greater cancer stages. This is intriguing as *mgl* has been recently shown to be the most highly transcribed of cysteine metabolizing genes^67^. Growth rate analysis of bacterial indicator species of CRC did not reveal statistically significant differences between disease states for any species analyzed, however this analysis was restricted by the limited presence of these species within healthy subjects. Future work that analyzes species that are present across all disease states is needed to understand if other keystone species play a role in CRC pathogenesis than determined previously by taxa abundance (Fig. S7, Table S9).

## DISCUSSION

There is compelling evidence implicating microbial H_2_S production as an environmental trigger of CRC, however ongoing studies investigating this link have been hampered by the field’s incomplete knowledge of sulfur metabolism within the human gut. Here, we performed a comprehensive characterization of the functional capacity of the human gut microbiome to conduct sulfur transformations and produce H_2_S. Overall, the data demonstrate that genes for microbial sulfur metabolism are more diverse than previously recognized, are widely distributed in the human gut microbiome, and are significantly associated with CRC.

Our investigation into the diversity and ecology of inorganic sulfur metabolism pathways observed that highly conserved functional genes encoding the final step of the sulfate reduction pathway — *dsrAB* — was harbored by six genera not typically targeted as SRB. Additionally, this investigation revealed that *asrABC*, an enzyme whose gene has homologous activity to Dsr, was both twice as abundant in total MAGs and was present in twice as many subjects than *dsrAB*. Together, these data highlight that genes for inorganic sulfur metabolism in the human gut are more widespread than previously established and that *asrABC* may be an important marker to measure the capacity of microbial sulfate reduction within cohorts. Since the diversity of microbial sulfatases have not been characterized, studies that characterize the catalytic efficiencies of these enzymes are needed to truly understand the implications of this expanded view of inorganic sulfur metabolism in the human gut.

While previous investigations of microbial H_2_S production and human disease have focused on SRB, bacteria that metabolize organic sulfur have been consistently associated with CRC risk. Prior to this work, *Bilophila wadsworthia* was the only bacterium known to produce H_2_S via taurine respiration. However, our analysis revealed *Desulfovibrio* spp. harbors genes for the characterized 4-step reductive pathway of taurine respiration, and an exploration of twelve taurine reduction pathways discovered two genera with genes for the complete 3-step reduction pathway. Further, 41 unique genera were found to have nearly complete 3- and 4-step pathways, revealing microbial targets for novel enzyme discovery or investigations of cooperative taurine metabolism.

Finally, analyses of diverse pathways for microbial cysteine and methionine degradation revealed that these sulfidogenic genes were distributed among diverse bacterial phyla and were abundantly present among subjects. Collectively, these analyses reveal that bacteria harboring pathways for organic sulfur metabolism are pervasive in the human gut and likely constitute the most abundant source of microbially derived H_2_S.

To determine whether sulfur metabolism was differentially associated along the colorectal carcinoma sequence (healthy → adenoma → carcinoma), presence of sulfidogenic genes was compared between healthy subjects, and patients with adenoma or carcinoma. Genes for both inorganic and organic sulfur metabolism were significantly associated with carcinoma, which is intriguing as both sulfate and taurine metabolism share a final metabolic step. These associations along with the pervasiveness of genes for organic sulfur metabolism in studied MAGs supports the unique hypothesis that organic sulfur metabolism by gut bacteria is a key mechanism linking a western diet and CRC risk (Fig. 5). A limitation of our study is that our analyses utilized MAGs which do not use all available metagenomic reads as a consequence of assembly and binning. Thus, it is possible that the abundance of sulfur genes present in the human gut is even greater than reported herein. However, the use of MAGs enabled taxonomic information, reconstruction of full pathways, and calculation of growth rates for selected CRC associated bacteria, thus providing robust new information regarding microbial sulfur metabolism in the human gut. Our findings serve as a foundation for future work characterizing the activity of sulfidogenic enzymes in diverse microbial species, exploring the expression of these genes as affected by health status, and examining microbial H_2_S induced tumorigenesis in animal models of CRC and human disease.

**Figure 5.**
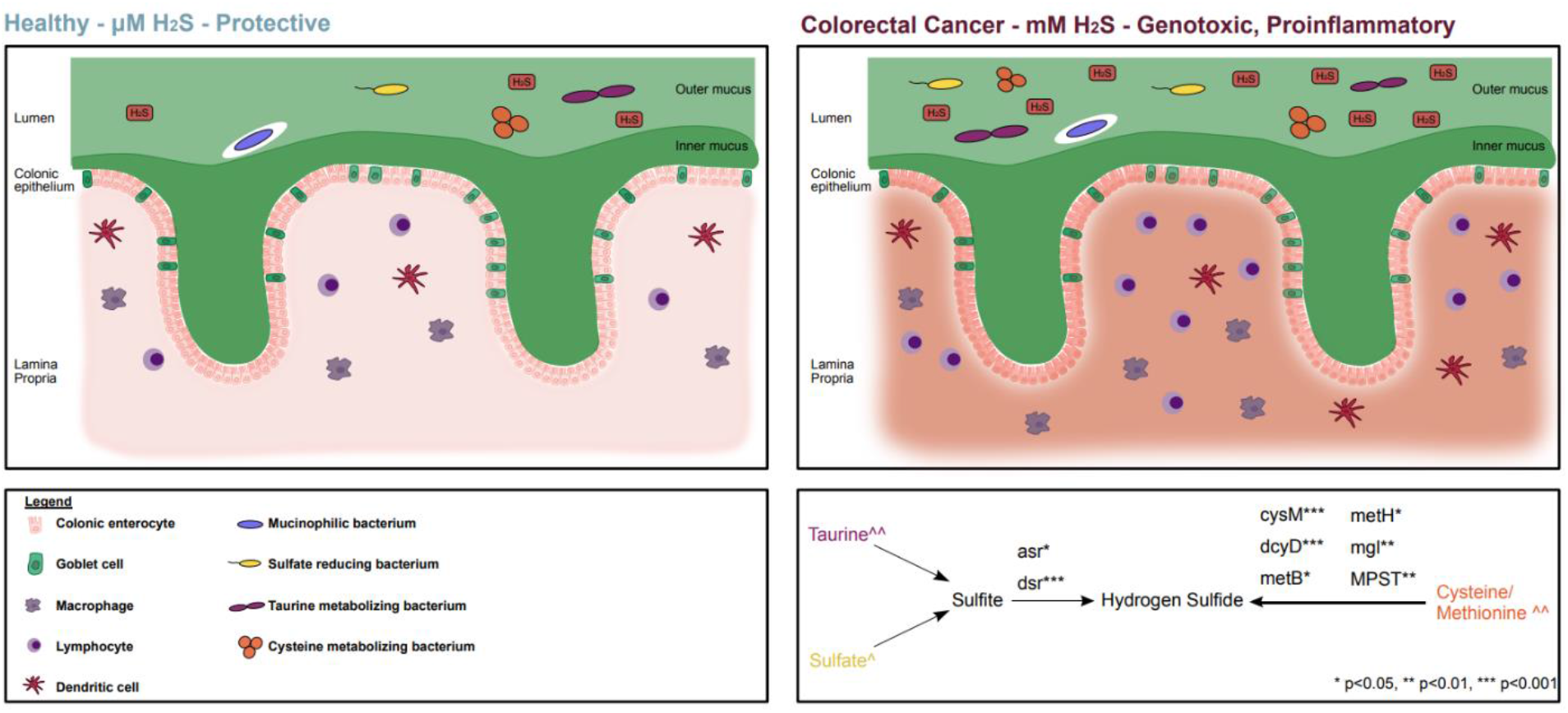
Organic sulfur metabolism by gut bacteria may be a key mechanism linking a western diet and CRC risk. The degradation of sulfomucins by mucolytic bacteria are a key source of inorganic sulfate for sulfate reducing bacteria. At μM concentrations, basal production of H_2_S through inorganic sulfate reduction exerts beneficial effects including gut barrier protection and fermentative hydrogen disposal. Intake of a western diet abundant in red and processed meat amplifies the production of taurine conjugated bile acids and increases colonic exposure to dietary sulfur amino acids (taurine, methionine, cysteine). In the context of a western diet, metabolism of organic sulfur amino acids by gut microbes drives the production of H_2_S to genotoxic and pro-inflammatory levels (mM concentration). Simplified pathways demonstrate genes for sulfur metabolism that were significantly associated with CRC. ^ indicates inorganic sulfur sources primarily provided by sulfated bile acids and sulfamucins. ^^ indicates organic sulfur sources provided by dietary sulfur amino acids and conjugated bile acids. Genes that have a non-random distribution across disease status as analyzed by chi-squared analysis are indicated by asterisks.

## METHODS

### Genomic survey of sulfidogenic genes in Human Microbiome Project genomes

An initial genomic survey was performed using 514 gastrointestinal genomes obtained from the Human Microbiome Project (HMP) in Fall 2018^68,69^. Reference sequences were obtained from the National Center for Biotechnology Information (NCBI) using searches for sulfidogenic genes from known residents of the human gut including “cysteine desulfhydrase”, “Cdl”, “Lcd”, “cystathionine-beta-synthase”, “L-methionine–gamma-lyase”, “dissimilatory sulfite reductase”, “dsrA”, “dsrB” and “dsrAB”. Searches of the HMP genes were then performed using BLAST (BLASTv2.8.1+)^70^, and alignments with greater than 60% identity and a minimum query coverage of 40 amino acids were retained. To filter non-homologous proteins, gene hits were compared to KEGG.

### Sulfur pathway visualization

Sulfur cycle reactions were created in ChemDraw Prime 16.0 and further modified in Affinity Designer.

### Downloading MAGs and accessing metadata

The previously reconstructed MAGs from the five cohorts were downloaded from https://opendata.lifebit.ai/table/sgb. The associated file that was downloaded, “download_files.sh” was used to download all 16,936 genomes from the 5 studies^35^. The link to download the metadata (https://www.dropbox.com/s/ht0uyvzzal6exs2/Nine_CRC_cohorts_taxon_profiles.tsv?dl=0) was found at http://segatalab.cibio.unitn.it/data/Thomas_et_al.html from two studies^71,72^ and filtered to only include the 5 cohorts used in our study. For samples that had AJCC TNM (Tumor, lymph Node, Metastasis) classification without a stage, the American Cancer Society guidelines to annotate stage based on TNM classification was used. For this study, all “high” (>90% complete, <5% contamination, <0.5% strain heterogeneity) and “medium” (>50% complete, <5% contamination) quality MAGs were included^36,73^.

### Gene annotation of MAGs

Prodigal module (prodigal version 2.6.3) of METABOLIC was used to run multiple threads with the -p meta option to annotate open reading frames (ORFs) on all MAGs^74,75^.

### Sulfur gene identification in MAG database

HMM - Hidden Markov Model (HMM) searches for protein sequences were performed using either hmm profiles from KEGG or custom profiles against a concatenated file of predicted ORFs from all 16,936 MAGs used for this study with HMMSearch version 3.3.0^76^, using trusted cut offs for all sulfur related sequences (Table S1). All HMMs used in this study are available at https://github.com/escowley/HumanGutBacterialSulfurCycle.

### Microbial sulfur pathway literature search

To determine the breadth of current knowledge regarding characterized H_2_S production in human gut bacteria and their association with CRC, a literature search was performed on genera revealed in our analysis. For MAGs that possessed sulfidogenic genes, the annotations for the “closest genus” were recorded. Web-searches were performed for each genera using the key words “colorectal cancer”, “sulfide”, and “H_2_S”. Genera that were previously associated with CRC or who have species that were characterized to produce H_2_S were noted in Supplemental Table 4.

### Taxonomic classifications of MAGs

Taxonomic classifications, “closest”designators, were determined previously by binning the MAGs with a reference set of genomes^36^. To verify taxonomic classifications for MAGs, a concatenated ribosomal protein tree, GTDB-Tk classification, and 16S rRNA alignment were done. Taxonomic classifications from the original study, GTDB-tk, and 16S can be found in Supplemental Table 6. 16S rRNA sequences were extracted from the MAGs using the ssu_finder function of checkM (version 1.0.11)^77^. From the 16,936 MAGs, 2531 16S rRNA sequences were extracted. Extracted 16S rRNA sequences were classified using with SINA aligner (version v1.2.11) (general options - bases remaining unaligned at the end should be attached to the last aligned base, reject sequences below 70% identity) using the search and classify feature (search and classify options - min identity with query sequence 0.95, number of neighbors per query sequence 10, sequence collection used - Ref-NR, search kmer candidates 1000, lca-quorum 0.8, search k-mer len 10, search kmer mm 0, search no fast, taxonomies used for classification SILVA, RDP, GTDB, LTP, EMBL-EBI/ENA), which classifies 16S sequences based on the least common ancestor and the SILVA reference database (release 138.1)^78–80^. Taxonomy was assigned using GTDB-Tk (version 1.3.0) with database release 95 with the classify_wf function^81–83^.

For the concatenated ribosomal protein tree, a curated Hidden Markov Models (HMM) database for single-copy ribosomal proteins (rpL2, rpL3, rpL4, rpL5, rpL6, rpL14, rpL14, rpL15, rpL16, rpL18, rpL22, rpL24, rpS3, rpS3, rpS8, rpS10, rpS17, rpS19)^84^ was used to identify these genes in all MAG using hmmsearch (version 3.3.0) using noise cutoffs (--cut_nc)^76^. Once identified, the protein sequences were extracted from the predicted ORFs and imported into Geneious Prime (v 11.1.5). Each sequence was aligned with a reference set using MAFFT (v 7.450, parameters Algorithm: Automatic, Scoring Matrix: BLOSUM62, Gap penalty: 1.53, Offset value: 0.123)^85^. The alignments were manually trimmed and a 95% gap masking threshold was applied to the resulting alignment. The resulting alignments were concatenated for each gene set. The concatenated alignments were exported in fasta format from Geneious and used as the input for making phylogenetic trees with IQTree using version 1.6.9 (-nt AUTO -m MFP -bb 1000 -redo -mset WAG,LG,JTT,Dayhoff -mrate E,I,G,I+G -mfreq FU -wbtl)^86^. The resulting tree file was imported into iToL for visualization and collapsing of nodes followed by final modifications in Affinity Designer^87^.

### Concatenated protein trees for dissimilatory sulfate reduction genes

To create the concatenated protein trees for *asrABC* and *dsrAB*, reference sequences for each set of genes from a diverse set of environments were utilized^47^. Hmmsearch (version 3.3.0) was used to identify asr and dsr sequences in the database of predicted ORFs^76^. The identified genes were extracted from the predicted ORFs and concatenation and tree building were done according to the protocol previously described for the ribosomal proteins.

### Summary calculations and statistical analysis for association of sulfur genes with disease state and stage of CRC

Summary calculations of total number of MAGs with the gene of interest per disease state, mean, median, and standard deviations were performed in R. Summary information can be found in Supplemental Table 3. To identify potential associations between presence of specific microbial sulfur genes and disease state, chi squared tests were performed. First, for each gene, each participant was binarized as either “presence” (at least 1 MAG with at least 1 copy of the gene of interest) or “absence” (no MAGs recovered from the participant’s sample had any copies of the gene of interest). For each gene, total presence and absence were tallied for each disease state (healthy, adenoma, carcinoma). Chi square tests were performed for each gene on a 2X3 matrix of presence/absence totals and disease states. P-value corrections were done using the Benjamini-Hochberg (BH) Procedure for final reported p-values. A significant p-value indicates the distribution of the gene among the disease states is not random. Summary values and statistics were all performed in R. Uncorrected and corrected p-values can be found in Supplemental Table 7. Dot plot visualizations with the proportion of participants in each of the 3 disease states with at least one copy of the gene in their MAGs normalized to total number of participants with that disease state as the size of the dot and mean number of MAGs with a copy of the gene per participant with at least 1 copy as the color of the dot were made using R with the cowplot package. Data used for these plots can be found in Supplemental Table 3.

Similarly, to identify associations between presence of specific microbial sulfur genes and colorectal cancer stage, we binarized the subset of studies with staging data as described above and performed binomial logistic regressions. Rao score tests were performed on the resulting binomial logistic regressions to evaluate for significance of each gene having evidence of association with staging of CRC. P-value corrections were done using the Benjamini-Hochberg (BH) Procedure for final reported p-values. Uncorrected and corrected p-values can be found in Supplemental Table 8. Presence/absence values for each gene in each stage were normalized to the total number of participants in that particular stage to generate distribution plots (Fig. S6).

### Metabolic reconstruction of MAGs

Metabolic reconstruction of each MAG was done using the METABOLIC-G program of METABOLIC (version 4.0)^75^. Summary information is available at https://github.com/escowley/HumanGutBacterialSulfurCycle.

### Determination of growth rates for colorectal cancer indicator bacteria

To determine the growth rates of the bacteria previously implicated as indicators of colorectal cancer in the gut community,^11,17^ we downloaded the original reads used to generate these genomes from the Hannigan, Yu, Zeller, and Feng studies. For the reads from the Hannigan study, we used the fasterq-dump (version 2.9.4) module of the SRA toolkit^88^, for the reads from the Yu and Zeller studies, we used the Aspera^89^ command line interface (CLI) ascp program (v3.9.1.168954), and for the reads from the Feng study, we used the Aspera CLI ascp program (v3.9.3.177167), respectively. For reads from the Zeller study, each participant has multiple read sets deposited, we chose the first listed paired-end read set for each genome listed in the European Nucleotide Archive (ENA) metadata. Reads from the Hannigan study were trimmed and quality filtered with metaWRAP (v1.2.2) using the read_qc^90^ module with the option “--skip-bmtagger”. We used Bowtie2 (v2.3.4.1)^91–93^ with the option “--reorder” to map MAGs classified as those bacteria to the original read sets and shrinksam^94^ version 0.9.0 with the “-u” flag to compress mapping files. We determined growth rates by generating un-filtered indexes of replication with iRep (version 1.10)^95^. We determined growth rates for the following organisms based on the closest taxonomic designator: *Fusobacterium mortiferum, Fusobacterium ulcerans, Fusobacterium nucleatum, Desulfovibrio piger, Bacteroides fragilis, Escherichia coli, Pyramidobacter piscolens, Clostridium difficile, Clostridium hylemonae, Porphyromonas asaccharolytica, Peptostreptococcus stomatis, Bilophila wadsworthia*, and *Odoribacter splanchnicus* (Table S9). Genomes from the species *F. nucleatum*, *P. piscolens*, and *O. splanchnicus* were not plotted. To determine differences in growth rates between CRC and healthy samples for all 3 studies, a Kruskal Wallis test was performed for each organism. To determine differences in growth rates across all disease states within a study for a particular organism, a Kruskal Wallis test was performed for each organism. P-value corrections were done using the Benjamini-Hochberg (BH) Procedure for final reported p-values. Uncorrected and corrected p-values can be found in Table S10. Code for statistical analysis and generation of plots can be found at https://github.com/escowley/HumanGutBacterialSulfurCycleLink.

## Supporting information

Supplementary Figures and References

## ACKNOWLEDGEMENTS

ESC is an MSTP student and was supported by an NLM training grant to the Computation and Informatics in Biology and Medicine Training Program (NLM 5T15LM007359) at UW-Madison, and in part by Medical Scientist Training Program grant T32GM008692. PGW was supported by the Cancer Education and Career Development Program grant T32CA057699. We thank the University of Wisconsin—Office of the Vice Chancellor for Research and Graduate Education and the University of Wisconsin-Department of Bacteriology for their support. JMR and HRG were supported by RO1CA204808. We (PGW and ESC) are thankful to the Anantharaman, Ridlon, and Gaskins lab for feedback and assistance throughout the process. We thank Madison Boissiere and Briawna Binion for their help with the literature search. We thank Michael Liou for statistical consulting and coaching throughout this project.

## AUTHOR CONTRIBUTIONS

PGW, ESC, HRG, and KA conceptualized the project and wrote the manuscript. PGW, SM, LL, PP performed microbial sulfur pathway literature search. AB downloaded the MAGs, ran Prodigal, and generated the growth rates and related figures. PGW and ESC ran HMM searches and generated figures. PGW, ESC, and KA analyzed data. ESC generated phylogenetic trees, generated metabolic reconstructions for MAGs, and performed statistical analyses.

## DATA AVAILABILITY STATEMENT

Raw reads for the original metagenomic studies are deposited at https://www.ebi.ac.uk/ena/browser/view/PRJEB12449, https://www.ebi.ac.uk/ena/browser/view/PRJEB7774, https://www.ebi.ac.uk/ena/browser/view/PRJEB10878, https://www.ncbi.nlm.nih.gov/bioproject/PRJNA389927/, and https://www.ebi.ac.uk/ena/browser/view/PRJEB6070. MAGs were constructed previously^35^ and were used for this study (see “downloading bins and accessing metadata” section of the methods). Code and HMM profiles are available at https://github.com/escowley/HumanGutBacterialSulfurCycleLink.

