## Supplementary Figures and References for "Diversity and distribution of sulfur metabolism in the human gut microbiome and its association with colorectal cancer"

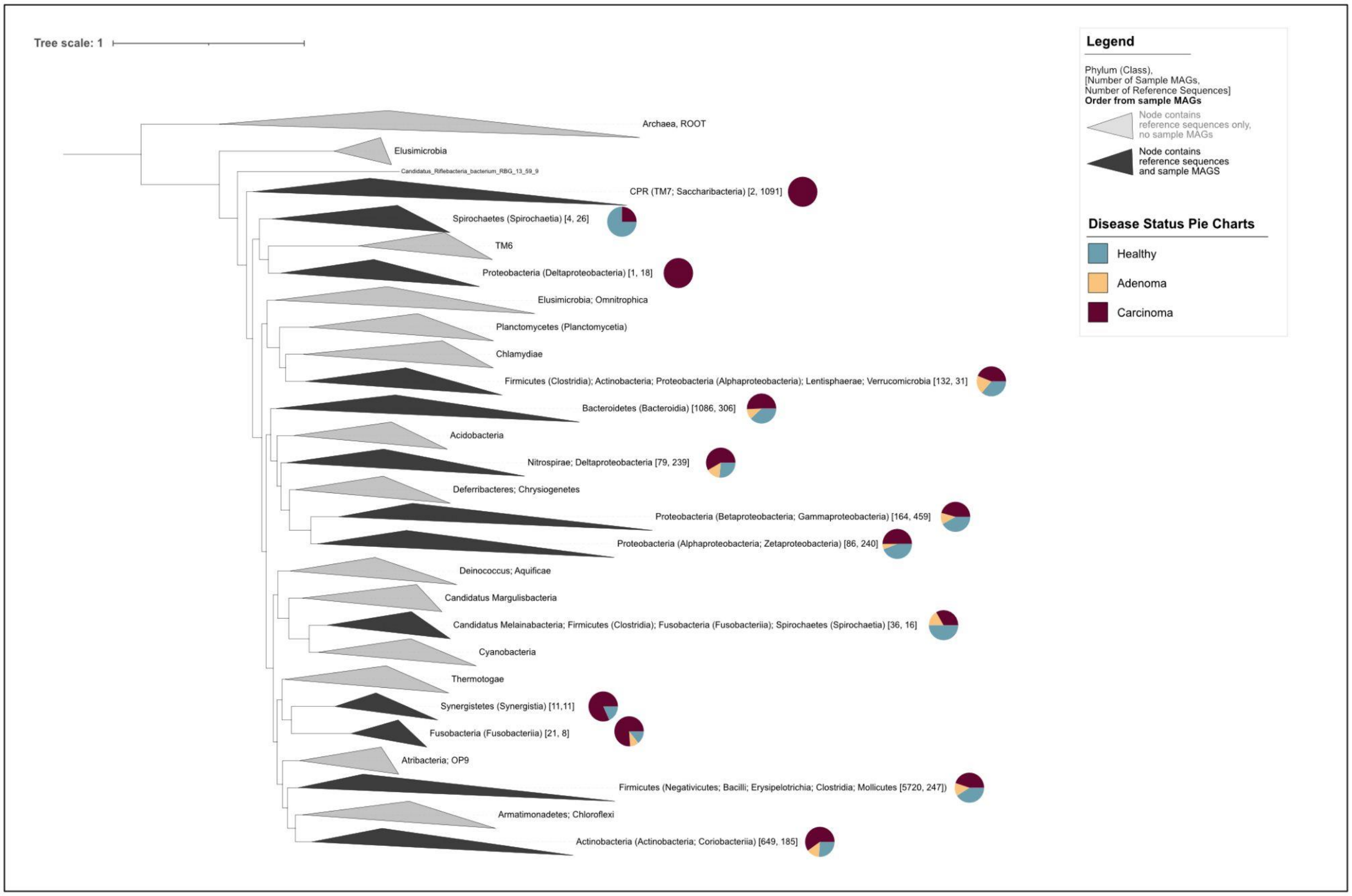


Supplemental Figure 1 - **Concatenated ribosomal protein tree of all samples demonstrating the full diversity of samples included in our analyses.** Concatenated protein tree of ribosomal proteins showing the diversity of bacterial MAGs included in our analyses. Gray clades only contain reference sequences, darker gray clades contain reference sequences and sequences from this study. Bracketed numbers indicate the sequence origin within each clade: [number of sequences from our study, number of sequences from references]. Pie charts indicate the disease state associated with sequences within each clade with blue indicating healthy, yellow adenoma, and maroon carcinoma.

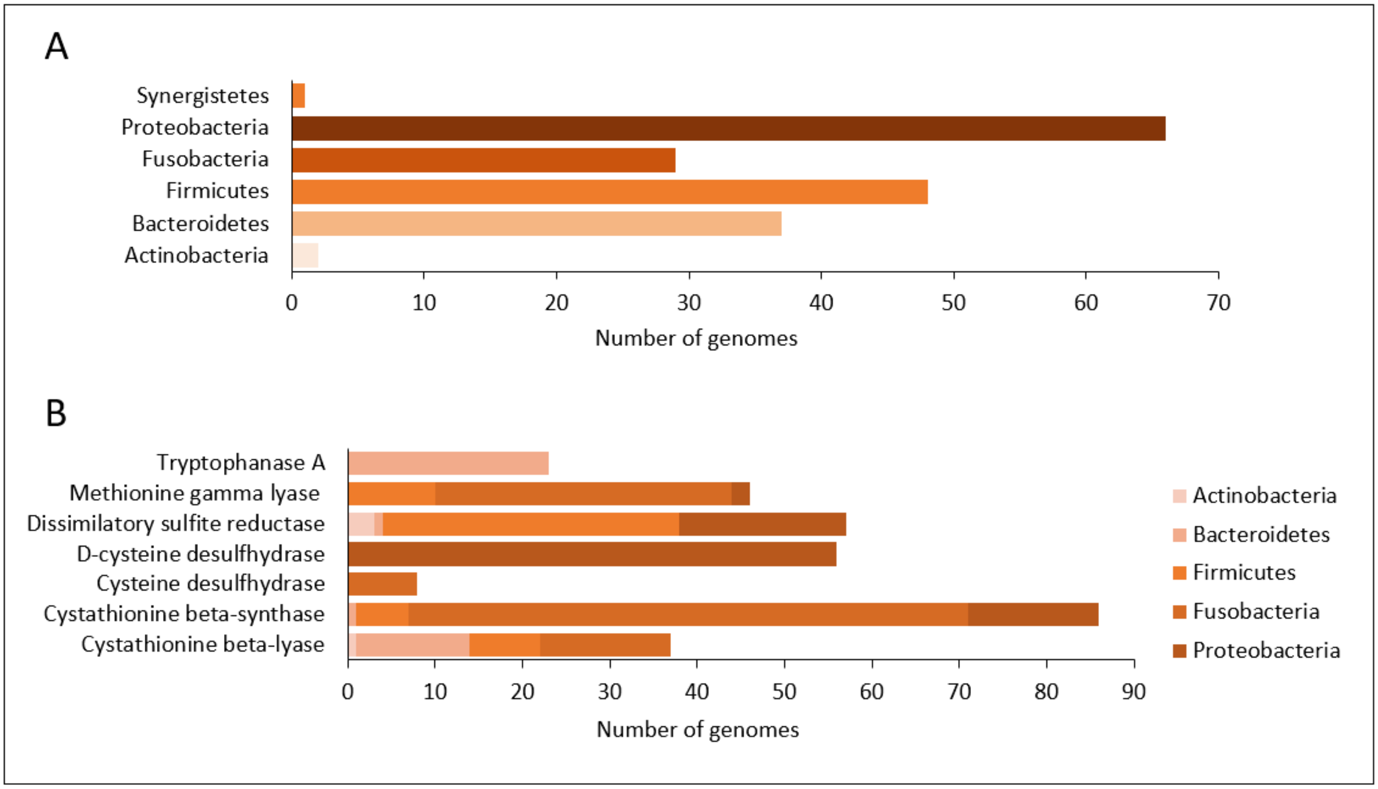


Supplemental Figure 2 - **Sulfidogenic functional genes are widespread in the human gut microbiome.** a. Identified sulfidogenic gastrointestinal genomes from the Human Microbiome Project distributed by phyla. b. Distribution of sulfidogenic genes from human gut genomes by phyla and functional gene class.

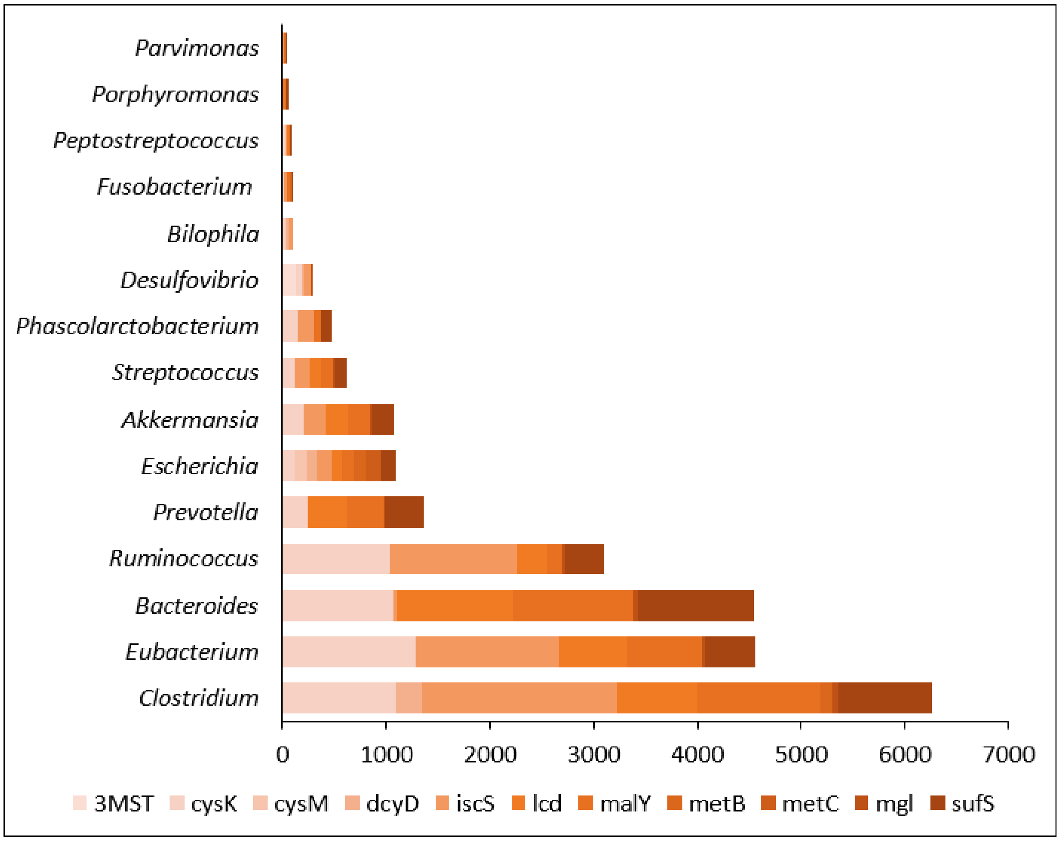


Supplemental Figure 3 - **Cysteine-metabolizing genes are abundant in bacteria commonly associated with CRC.** Results of a phylogenetic analysis of predicted cysteine-producing genes from MAGs. Displayed are the number of genomes containing cysteine-metabolizing genes in selected bacteria commonly associated with CRC.

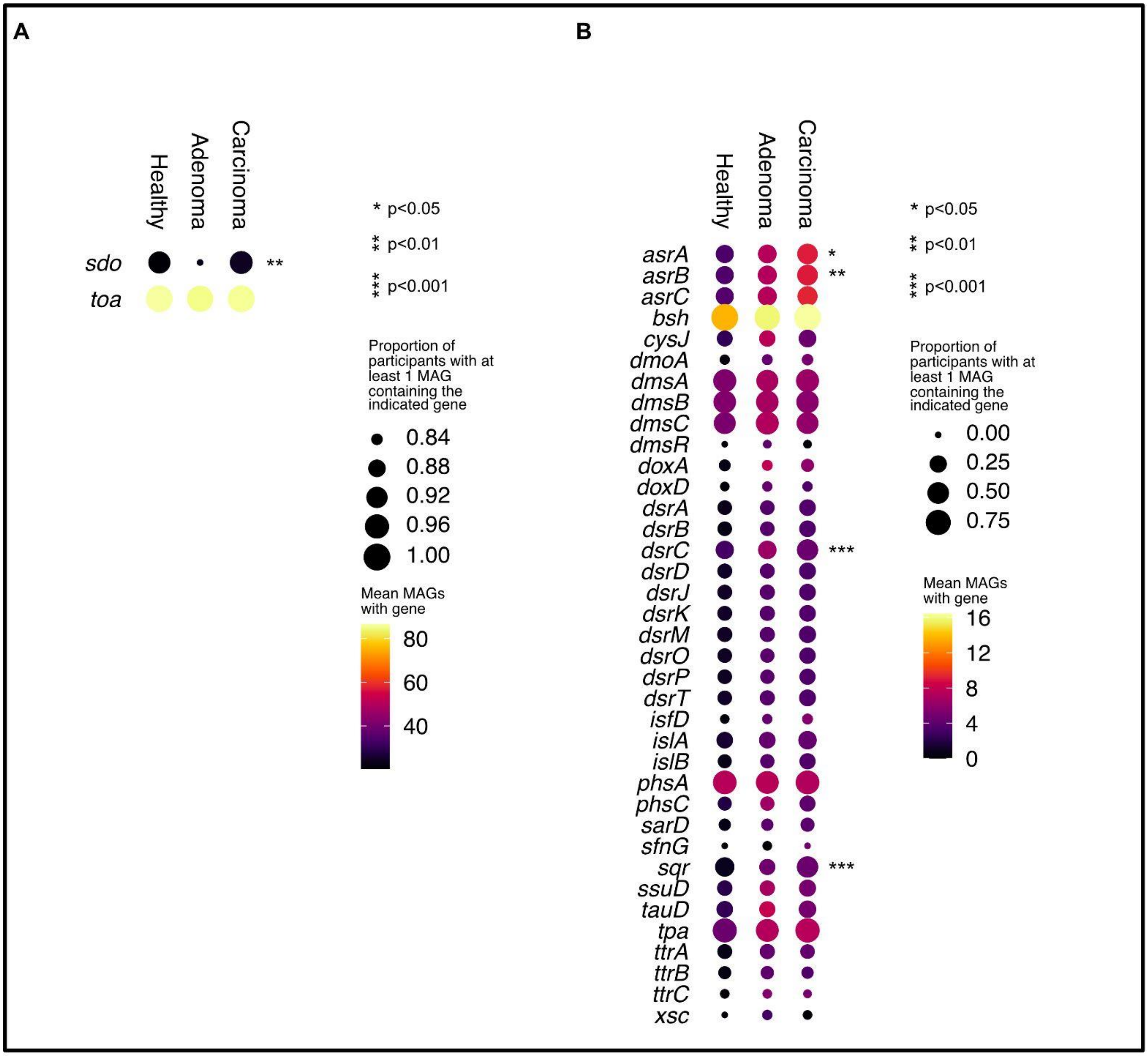


Supplemental Figure 4 - **Comprehensive dot plots of (A) highly-abundant and (B) all other genes analyzed related to bacterial taurine and sulfur metabolism across three disease states: healthy, adenoma, and carcinoma.** The size of each dot indicates the proportion of participants in each disease state with at least 1 copy of the indicated gene in their bacterial MAGs and the color of each dot indicates the mean number of MAGs with that gene in the subset of participants that have at least 1 copy of the gene. Genes that have a non-random distribution across disease status as analyzed by chi-squared analysis are indicated by asterisks.

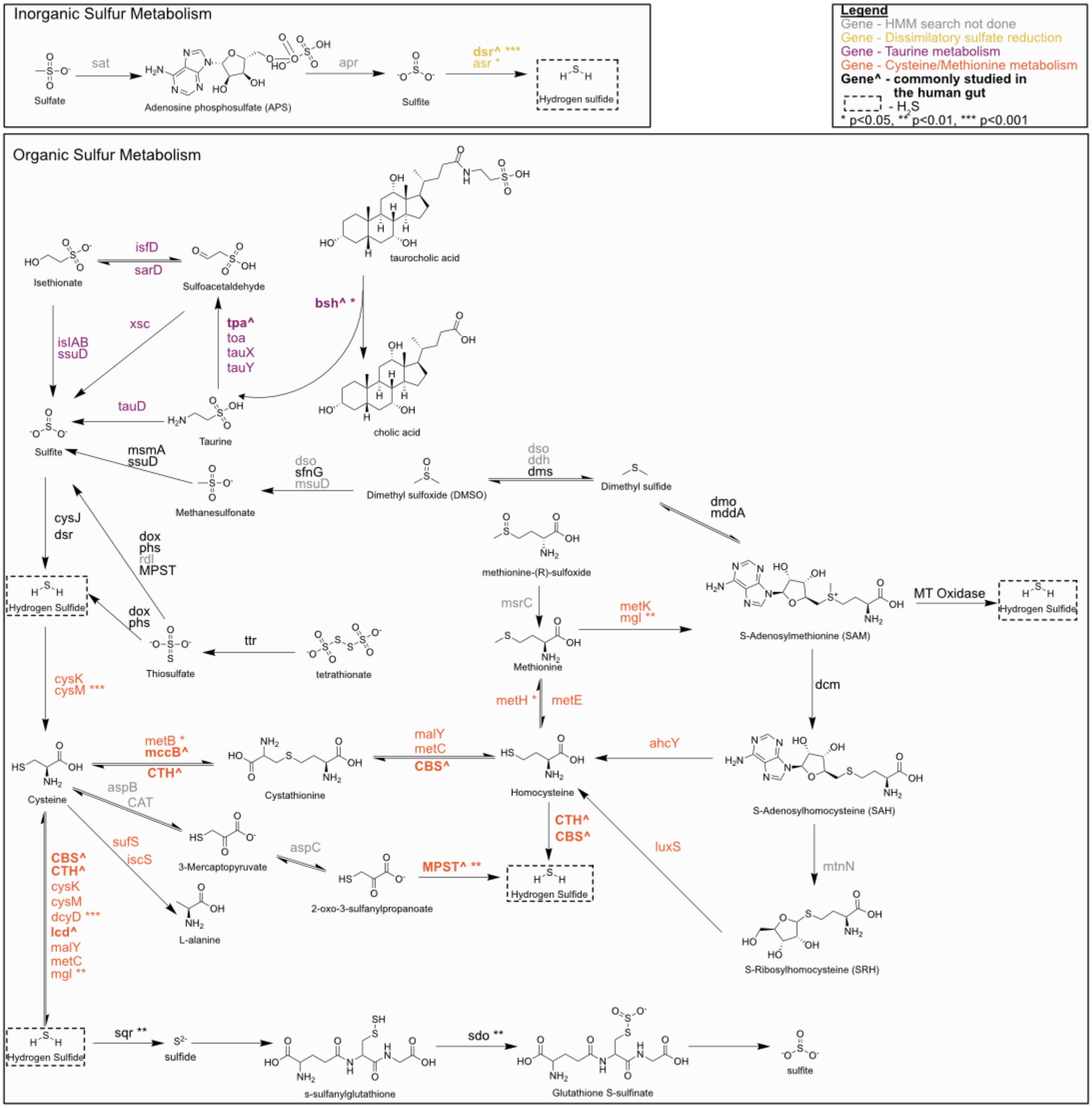


Supplemental Figure 5 - **Sulfidogenic pathways are significantly associated with colorectal cancer.** Bacterial sulfur cycling results in the production of genotoxic $H_2S$ (dashed box) via metabolism of inorganic sulfate (yellow) or organic sulfur amino acids like cysteine and methionine (maroon) or taurine (orange). Previous studies of bacterial sulfidogenesis in the human gut have focused mainly on genes harbored by *Bilophila*, *Fusobacterium*, and the sulfate reducing bacteria (bolded with a “^”). Genes listed were analyzed by this study except those listed in gray. Genes that were found to be significantly different between disease states as described in the main text are indicated with asterisks. Reactions are not balanced and only the main sulfur component reactants and products are shown. Some intermediate steps are not shown.

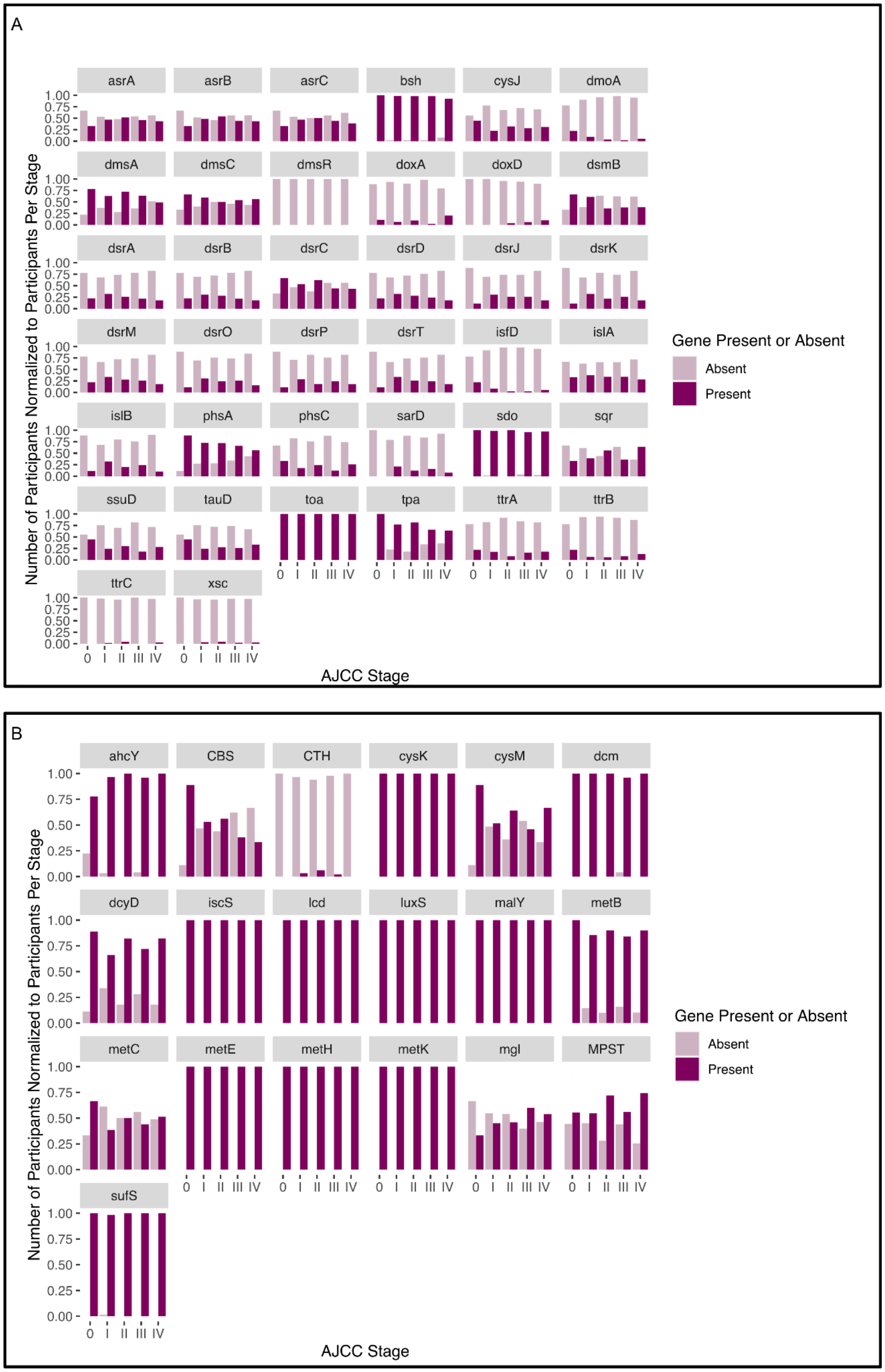


Supplemental Figure 6 - **Genes for bacterial sulfur metabolism are differentially distributed across different stages of colorectal cancer.** Distributions of genes related to bacterial taurine and sulfur metabolism (A) and cysteine and methionine metabolism (B). The number of participants with the particular gene present or absent is normalized to the total number of participants in each specified stage.

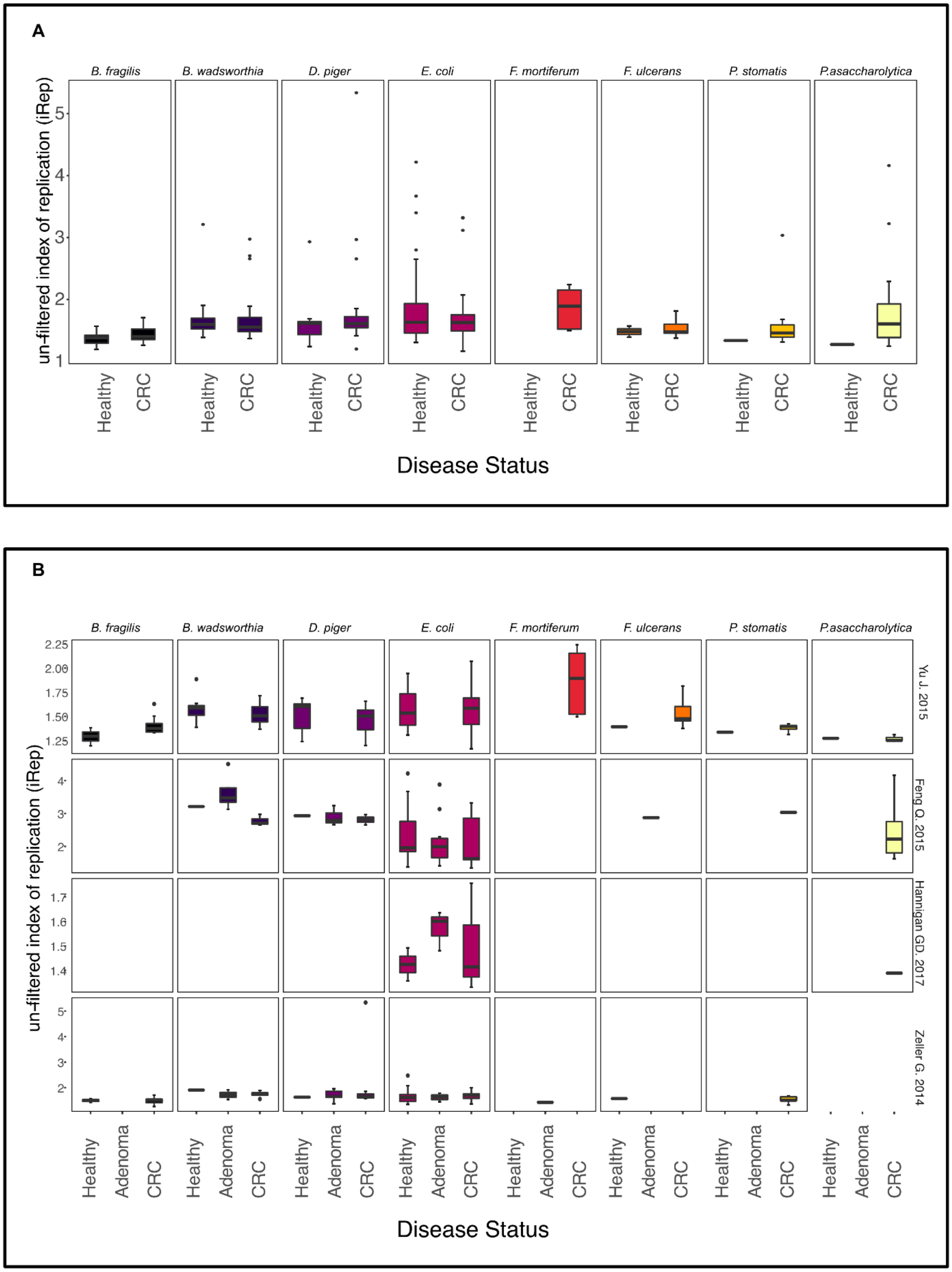


Supplemental Figure 7- **Indicator species for CRC are more likely to be present in participants with CRC and there are no differences in growth rates across disease states for any organism analyzed.** Un-filtered iRep growth rates, an estimation of instantaneous bacterial population growth based on a comparison of genome coverage at the origin and terminus of replication, were generated (values before filtering by iRep default thresholds) for genomes from the three studies collectively, representing both healthy and CRC samples (A) and of genomes separated by individual study within the healthy, adenoma, and CRC samples (B).
